# Carbendazim-resistance associated β_2_-tubulin substitutions increase deoxynivalenol biosynthesis by reducing the interaction between β_2_-tubulin and IDH3 in *Fusarium graminearum*

**DOI:** 10.1101/760595

**Authors:** Zehua Zhou, Yabing Duan, Mingguo Zhou

## Abstract

Microtubule is a well-known structural protein participating in cell division, motility and vesicle traffic. In this study, we found that β_2_-tubulin, one of the microtubule components, plays an important role in regulating secondary metabolite deoxynivalenol (DON) biosynthesis in *Fusarium graminearum* by interacting with isocitrate dehydrogenase subunit 3 (IDH3). We found IDH3 negatively regulate DON biosynthesis by reducing acetyl-CoA accumulation in *F. graminearum* and DON biosynthesis was stimulated by exogenous acetyl-CoA. In addition, the expression of IDH3 significantly decreased in the carbendazim-resistant mutant nt167 (Fgβ_2_ ^F167Y^). Furthermore, we found that carbendazim-resistance associated β_2_-tubulin substitutions reducing the interaction intensity between β_2_-tubulin and IDH3. Interestingly, we demonstrated that β_2_-tubulin inhibitor carbendazim can disrupt the interaction between β_2_-tubulin and IDH3. The decreased interaction intensity between β_2_-tubulin and IDH3 resulted in the decreased expression of IDH3, which can cause the accumulation of acetyl-CoA, precursor of DON biosynthesis in *F. graminearum*. Thus, we revealed that carbendazim-resistance associated β_2_-tubulin substitutions or carbendazim treatment increases DON biosynthesis by reducing the interaction between β_2_-tubulin and IDH3 in *F. graminearum*. Taken together, the novel findings give the new perspectives of β_2_-tubulin in regulating secondary metabolism in phytopathogenic fungi.

**Author Summary:** The deoxynivalenol (DON) biosynthesis is increased in carbendazim-resistant strains in *Fusarium graminearum*. To date, the molecular mechanism between the carbendazim-resistant substitution and the increased DON production remained elusive. Here we found that acetyl-CoA-associated enzyme IDH3 negatively regulates acetyl-CoA and DON biosynthesis. Moreover, β_2_ tubulin interacted with IDH3 physically and increase its expression. We further found that carbendazim-resistant substitution in β_2_ tubulin reducing the interaction between β_2_ tubulin and IDH3, which resulted in the decreased expression of IDH3. In addition, we demonstrated that carbendazim disrupting the binding between β_2_ tubulin and IDH3, which also decreases the expression of IDH3. Taken together, our results give a newly insights into the mechanism of β_2_ tubulin and its carbendazim-resistant substitution in regulating DON biosynthesis.

## Introduction

Fusarium head blight (FHB) caused by the filamentous fungus *Fusarium graminearum* is a devastating fungal disease in wheat and other small grain cereals all over the world [1]. The disease can not only lead to loss in yield and quality, but also a severe threat to human and animal health due to the mycotoxins in infested grains [1,2]. Among the various mycotoxins found in small grains, deoxynivalenol (DON) has been shown to be the most common mycotoxin contaminant associated with FHB [1]. In addition, as a potent inhibitor of protein synthesis, DON is carcinogenic to man and mammals [3,4]. In the past decades, all the *TRI* genes involved in trichothecene biosynthesis have been well-characterized [1,5]. In *F. graminearum*, trichothecenes are synthesized from farnesyl pyrophosphate (FPP), which is synthesized via the isoprenoid biosynthetic pathway [6]. Mevalonate, the key intermediate of this the pathway, is produced from three acetyl-CoA molecules. As a central molecule in cell metabolism, acetyl-CoA can be generated in the mitochondrial matrix, cytosol and peroxisome [7]. The research findings in metabolic engineering indicated that by modulation the production and consumption pathway of intracellular acetyl-CoA can directly increase the production of secondary metabolites, which showed that the production of acetyl-CoA is important for secondary metabolism [8–10].

As a key molecule, acetyl-CoA is predominantly generated by the tricarboxylic acid (TCA) cycle in the mitochondrial matrix [7]. In addition, isocitrate dehydrogenase (IDH) is one of the regulatory enzymes of the TCA cycle [11,12]. The researches of IDH focused on both eukaryotes and prokaryotes. Despite the fact that this enzyme converts isocitrate to 2-oxoglutarate and supports the TCA cycle, it is divided divide into cytosolic type and mitochondrial type in all eukaryotic organisms [11]. In HepG2 cells, silencing of cytosolic *IDH1* increases cell vulnerability by damaging the cellular redox status [13]. A similar phenomenon was also observed in melanoma cells and mice [14,15]. In *Aspergillus niger* WU-2223L, over-expression of mitochondrial *icdA* significantly decreased citric acid production [16].

Currently, several strategies have been developed to control this disease as well as reduce mycotoxin contamination in grains, of which chemical control is the major approach for controlling FHB. In China, carbendazim and other benzimidazole fungicides have been widely used to control the disease [17]. However, the emergence of resistant pathogen populations in the field has made carbendazim less effective since the 1990s [18]. Our previous researches showed that the amino acid substitutions of β_2_-tubulin at codons 167, 198 or 200 are responsible for carbendazim resistance and the substitution at codon 167 is dominant, occupying more than 90% [19–21]. Furthermore, carbendazim resistance is positively correlated with the DON biosynthesis ability of *F. graminearum* [22].

Numerous research findings have revealed that the microtubules (MTs) cytoskeleton play an important role in regulating metabolic processes by interacting with cytoplasmic enzymes, mitochondrial outer membrane and hypoxia inducible factor (HIF)-1 [23]. MTs can bind almost all glycolytic enzymes to some degree and change their activity. For instance, phosphofructokinase (PFK) and triosephosphate isomerase (TPI) exhibited reduced activity after binding to MTs, hexokinase (HK) and pyruvate kinase (PK) showed greater activity after binding to MTs [24–27]. Our transcriptome data indicated that mutation at codon 167 of β_2_-tubulin had a global effect on the expression of metabolic enzymes (enzymes present in the cytoplasm and mitochondria), such as isocitrate dehydrogenase, one of the regulatory enzymes in the TCA cycle (our unpublished data). However, the underlying molecular mechanism of the carbendazim-resistant substitutions of β_2_-tubulin in regulating DON biosynthesis remains elusive.

The aim of the present was to reveal the underlying molecular mechanism of carbendazim-resistance associated β_2_-tubulin substitutions in increasing of DON biosynthesis. The results showed that carbendazim-resistant substitutions in β_2_-tubulin could reduce the expression of IDH3 (a subunit of the regulatory enzyme in the TCA cycle) by reducing the binding between β_2_-tubulin and IDH3, causing the accumulation of cytosolic acetyl-CoA and then accelerating DON biosynthesis in *F. graminearum*.

## Results

### Substitution of β2-tubulin conferring carbendazim resistance decreases isocitrate dehydrogenase expression in F. graminearum

Our previous studies suggested that the substitution of β_2_-tubulin F167Y, conferring carbendazim resistance, exhibited significantly increased DON biosynthesis [22]. To elaborate the phenomenon, we analyzed the transcriptome difference between the wild-type strain 2021 and the F167Y mutant nt167. The results revealed that the F167Y substitution of β_2_-tubulin had a global effect on various pathways, including primary and secondary metabolism as compared with the wild-type strain 2021 (our unpublished data). In addition, qRT-PCR demonstrated that the transcriptional level of isocitrate dehydrogenase, a regulatory enzyme involved in the TCA cycle, dramatically decreased in nt167 (Fig 1A). Furthermore, we compared the intracellular acetyl-CoA production, which is metabolized in the TCA cycle, between 2021 and nt167. The results showed that the intracellular acetyl-CoA increased by 15% in nt167 as compared with 2021 (Fig 1B). Our findings also showed that more bulbous structures were observed in nt167 in comparison to 2021 (Fig 1C), which are associated with DON biosynthesis in *F. graminearum* [28,29].

**Fig 1.**
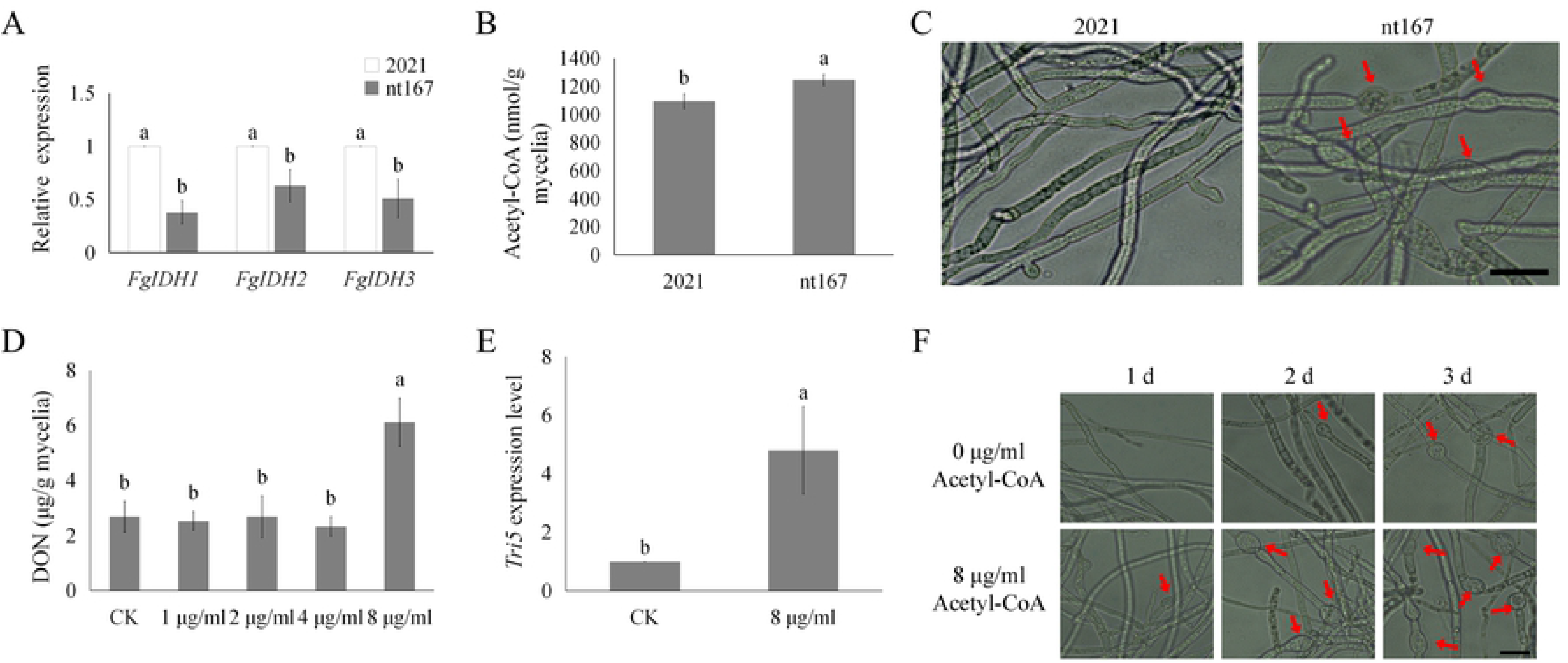
F167Y substitution reduces FgIDH3 expression and subsequently increases acetyl-CoA concentration in *F. graminearum*. A, Relative expression levels of *FgIDH1*, *FgIDH2* and *FgIDH3* between the wild-type strain 2021 and the mutant nt167. The relative expression level in 2021 was set to 1. Error bar indicates standard deviation (SD) calculated from the data of three replicates. B, Acetyl-CoA production in 3-day-old YEPD cultures of the wild-type strain 2021 and the mutant nt167. Bars denote standard errors from three repeated experiments. Values on the bars followed by the same letter are not significantly different at P = 0.05, according to Fisher’s LSD test. C, 3-day-old hypha of the wild-type strain 2021 and the mutant nt167 in GYEP cultures were examined for bulbous structures (labelled with arrows). Bar = 20 μm. D, DON production in GYEP cultures of 2021 treatment with 0, 1, 2, 4 and 8 μg/ml acetyl-CoA after incubation for 7 days. Error bar indicates standard deviation (SD) calculated from the data of three replicates. E, *Tri5* expression was assayed by qRT-PCR with RNA samples isolated from 2021, cultures were incubated in the presence/absence of 8 μg/ml acetyl-CoA for 4 days. Acetyl-CoA was added after a 3-day incubation. The relative expression level of *Tri5* in cultures without exogenous acetyl-CoA was set to 1. Error bar indicates standard deviation (SD) calculated from the data of three replicates. F, GYEP cultures of 2021 treated with or without 8 μg/ml acetyl-CoA were examined for bulbous structures (marked with arrows) after acetyl-CoA was added for 1, 2 and 3 days. Bar = 20 μm.

### Exogenous acetyl-CoA stimulates DON biosynthesis in *F. graminearum*

Previous studies reported that the concentration of acetyl-CoA is positively related to DON biosynthesis in *F. graminearum*, but the effect of exogenous acetyl-CoA on DON biosynthesis remain unknown [30]. In our study, the wild-type strain 2021 was treated with 1, 2, 4 or 8 μg/ml acetyl-CoA after incubation for 3 days in GYEP and assayed for DON production after incubation for another 4 days. The results showed that DON biosynthesis significantly increased in cultures treated with 8 μg/ml acetyl-CoA in 2021 (Fig 1D). When assayed by qRT-PCR, the expression of *Tri5* significantly increased in cultures treated with 8 μg/ml acetyl-CoA in 2021 (Fig 1E). In addition, when cultures were treated with 8 μg/ml acetyl-CoA, more abundant bulbous structures were observed compared to the control group (Fig 1F). Moreover, the Fghxk deletion mutant ΔFghxk, recorded similar results after treatment with exogenous acetyl-CoA (S1 Fig). An earlier study demonstrated that *TRI* genes transcription was significantly observed in GYEP, after incubation for 3 days [22]. Our findings showed that exogenous acetyl-CoA did not affect DON biosynthesis prior to the high transcription of *TRI* genes (S2 Fig). These results indicated that acetyl-CoA treatment stimulated DON biosynthesis and cellular differentiation after *TRI* genes transcription.

### Sequence analysis and subcellular localization reveals the functional differentiation of FgIDHs

There are two types of IDHs in a eukaryotic cell: 1) mitochondrial type and 2) cytosolic type [11]. Although these enzymes catalyze the same reaction, we hypothesize that there is functional differentiation between both types of enzymes. To gain a deeper insight into IDH functional differentiation in fungi, we conducted a phylogenetic analysis of three IDH enzymes, which were contained in nine of the most common phytopathogenic fungi, including *F. graminearum.* The results showed that the IDH enzymes are significantly divided into three groups (Fig 2A). The phylogenetic tree indicated that IDH enzymes might have different biological functions in *F. graminearum.* To gain a better understanding of the biological functions of three FgIDHs, we identified their subcellular localization in *F. graminearum*. We constructed ectopic transformants of each FgIDH-GFP fusion protein with the native promoter region and ORF (excluding the stop codon). All transformants were verified by western blotting assay and observed by confocal microscopy of the FgIDHs-GFP. The FgIDH1-GFP strain stained by CMAC, a dye that labels the vacuole, was localized in the vacuoles (Fig 2B). FgIDH2-GFP and FgIDH3-GFP strains, stained by Mito Tracker Red, a dye that labels the mitochondria [31], showed a colocalization of the red fluorescent probe with FgIDH2-GFP and FgIDH3-GFP signals in the vegetative hyphae, indicating that FgIDH2-GFP and FgIDH3-GFP are localized in the mitochondria of *F. graminearum* (Fig 2B). These results suggested that FgIDH2 and FgIDH3 might have similar biological functions while FgIDH1 might play a different role in *F. graminearum*.

**Fig 2.**
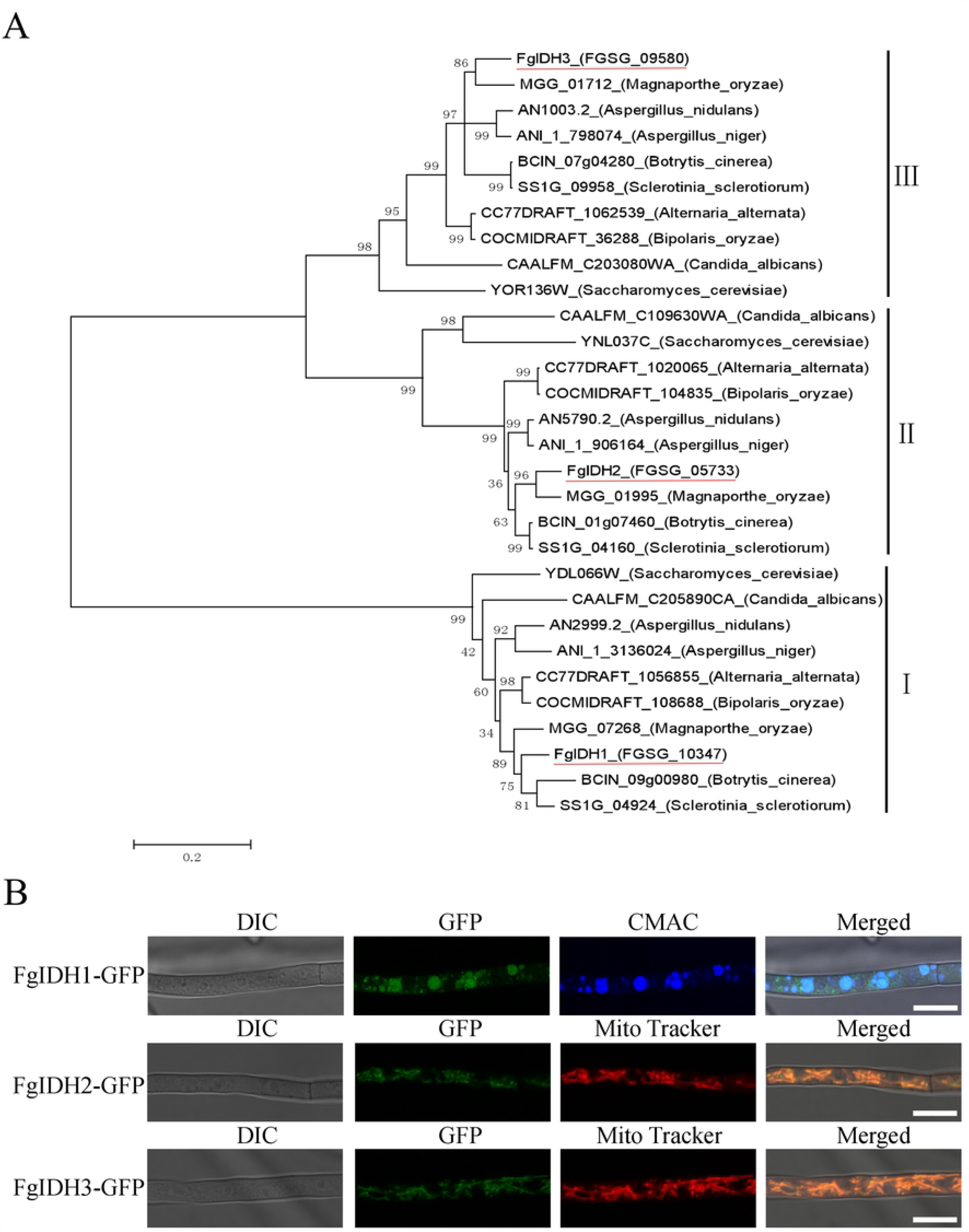
Phylogenetic analysis and subcellular localization of FgIDH in *F. graminearum*. A, Phylogenetic analysis of the putative IDH proteins from nine pathogenic fungi and *F. graminearum*. The amino acid sequences of each IDH orthologue was aligned using the neighbor-joining method with Mega 5.0 software. B, Subcellular localization of FgIDHs in *F. graminearum*. Localization of FgIDH1-GFP to the vacuolar in vegetative mycelia. Localization of FgIDH2-GFP and FgIDH3-GFP to the mitochondria in vegetative mycelia.

### Deletion of FgIDH3 significantly increases DON biosynthesis and TRI5 gene expression

Given that the substitution of β_2_-tubulin F167Y significantly decreased the expression of *FgIDHs* and the functional divergence of three FgIDHs, we generated the deletion mutants ΔFgIDH1, ΔFgIDH2 and ΔFgIDH3 from the wild-type strain 2021 by the split-marker approach to investigate their roles in DON biosynthesis in *F. graminearum* (S3-S5 Fig). As indicated, aerial mycelia on PDA plates were reduced for ΔFgIDH2 and ΔFgIDH3 but not for ΔFgIDH1 (Fig 3A and S3 Table). Hyphal morphology did not differ between the *FgIDH* deletion mutants and 2021 (S6 Fig).

**Fig 3.**
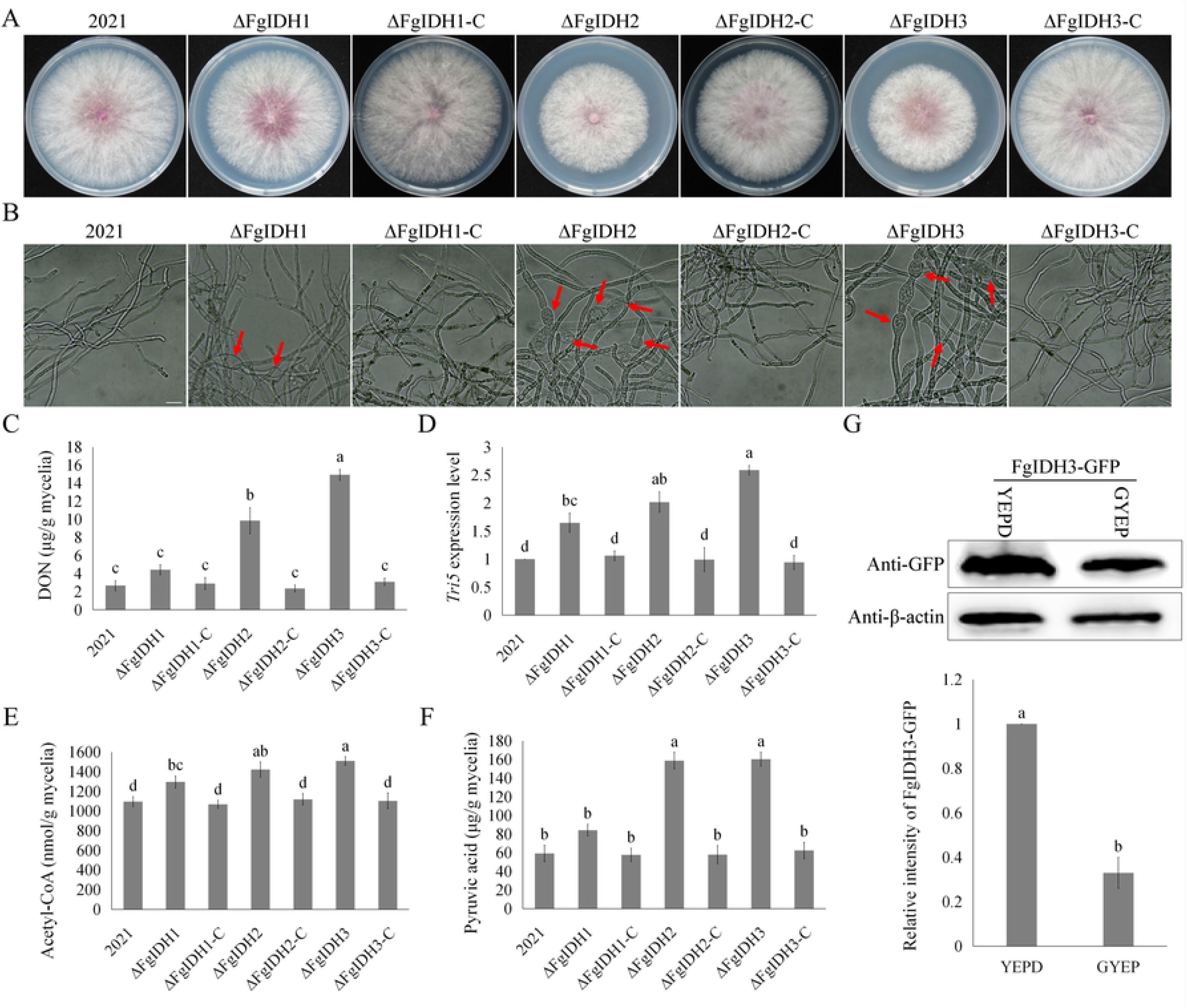
FgIDH3 negatively regulates DON biosynthesis and *TRI* gene expression. A, The wild-type strain 2021 and each IDH mutant was grown on PDA at 25°C for 3 days. B, 3-day-old GYEP cultures of the wild-type strain 2021 and each mutant was examined for bulbous structures (labelled with arrows). Bar = 20 μm. C and D, DON production and *Tri5* expression in GYEP cultures of the wild-type strain 2021 and each mutant. Error bar indicates standard deviation (SD) calculated from the data of three replicates. E and F, Acetyl-CoA and pyruvic acid production in 3-day-old YEPD cultures of the wild-type strain 2021 and each mutant. Error bar indicates standard deviation (SD) calculated from the data of three replicates. G, The expression of FgIDH3-GFP protein was assayed from 3-day-old YEPD (set to 1) and GYEP cultures, determined by western blotting assay with the anti-GFP antibody. The protein sample also incubated with the anti-β-actin antibody as a reference.

When assayed with GYEP cultures, an increase was found in DON biosynthesis by over 3.7- and 5.6-folds in ΔFgIDH2 and ΔFgIDH3, respectively (Fig 3C). In addition, *Tri5* expression in ΔFgIDH2 and ΔFgIDH3 was increased by 2.6- and 2.0-folds in comparison to 2021, respectively (Fig 3D). These data suggested that *FgIDH2* and *FgIDH3* but not *FgIDH1* play more important but negative roles in the regulation of DON biosynthesis. Furthermore, we assayed DON production with wheat grains. Similar results were obtained with GYEP cultures, DON production was 3.4- and 4.4-folds higher in wheat grains of ΔFgIDH2 and ΔFgIDH3 than those of 2021, respectively (S3 Table). We further determined acetyl-CoA and pyruvic acid production in each mutant. As expected, the acetyl-CoA production increased by a factor of 18.2, 29.7 and 37.7% in ΔFgIDH1, ΔFgIDH2 and ΔFgIDH3 compared to 2021, respectively (Fig 3E). Similarly, pyruvic acid production was 1.4-, 2.7- and 2.7-folds higher in comparison with 2021, respectively (Fig 3F).

When assayed for FgIDH3 expression in YEPD and GYEP cultures, we found that the translational level of FgIDH3 was down-regulated under DON-producing conditions, which are consistent with its role in DON regulation (Fig 3G). After being incubated in GYEP cultures for 3 days, swollen hyphal structures were rarely observed in 2021, but abundant swollen hyphal structures were found in ΔFgIDH2 and ΔFgIDH3 (Fig 3B). These phenomena are consistent with the significantly increased DON biosynthesis in each mutant.

Taken together, these results indicated that *FgIDH2* and *FgIDH3* play a major role in regulating DON biosynthesis by increasing the intracellular production of acetyl-CoA. More importantly, these phenomena from our genetic manipulations are consistent with the metabolic change of nt167, which resulted from the F167Y mutation of β_2_-tubulin.

### The increased DON biosynthesis ability interrelates with carbendazim-resistant substitutions of β2-tubulin

To test whether or not the carbendazim-resistant and non-carbendazim-resistant substitutions of β_2_-tubulin or substitutions of other microtubules resulted in increased DON biosynthesis in *F. graminearum*, we examined DON production and *Tri5* expression in several carbendazim-resistant strains. As shown in Fig 4A and 4B, all carbendazim-resistant mutants (Fgβ_2_^Y50C^, Fgβ_2_^E198Q^, Fgβ_2_^E198L^, Fgβ_2_^E198K^ and Fgβ_2_^F200Y^) increased DON production and *Tri5* expression while non-carbendazim-resistant substitution mutants (Fgβ_2_^F240L^ and Fgβ_1_^F167Y^) exhibited similar DON production and *Tri5* expression in comparison to the wild-type strain 2021. Furthermore, we determined the intracellular production of acetyl-CoA and pyruvic acid in these mutants. As expected, there was an increase in acetyl-CoA production in all the carbendazim-resistant strains in comparison to 2021, as well as pyruvic acid production. However, there was no observed difference of acetyl-CoA and pyruvic acid production between non-carbendazim-resistant strains and 2021 (Fig 4C and 4D). In summary, the substitutions of β_2_-tubulin conferring carbendazim-resistance resulted in increased DON biosynthesis in *F. graminearum*.

**Fig 4.**
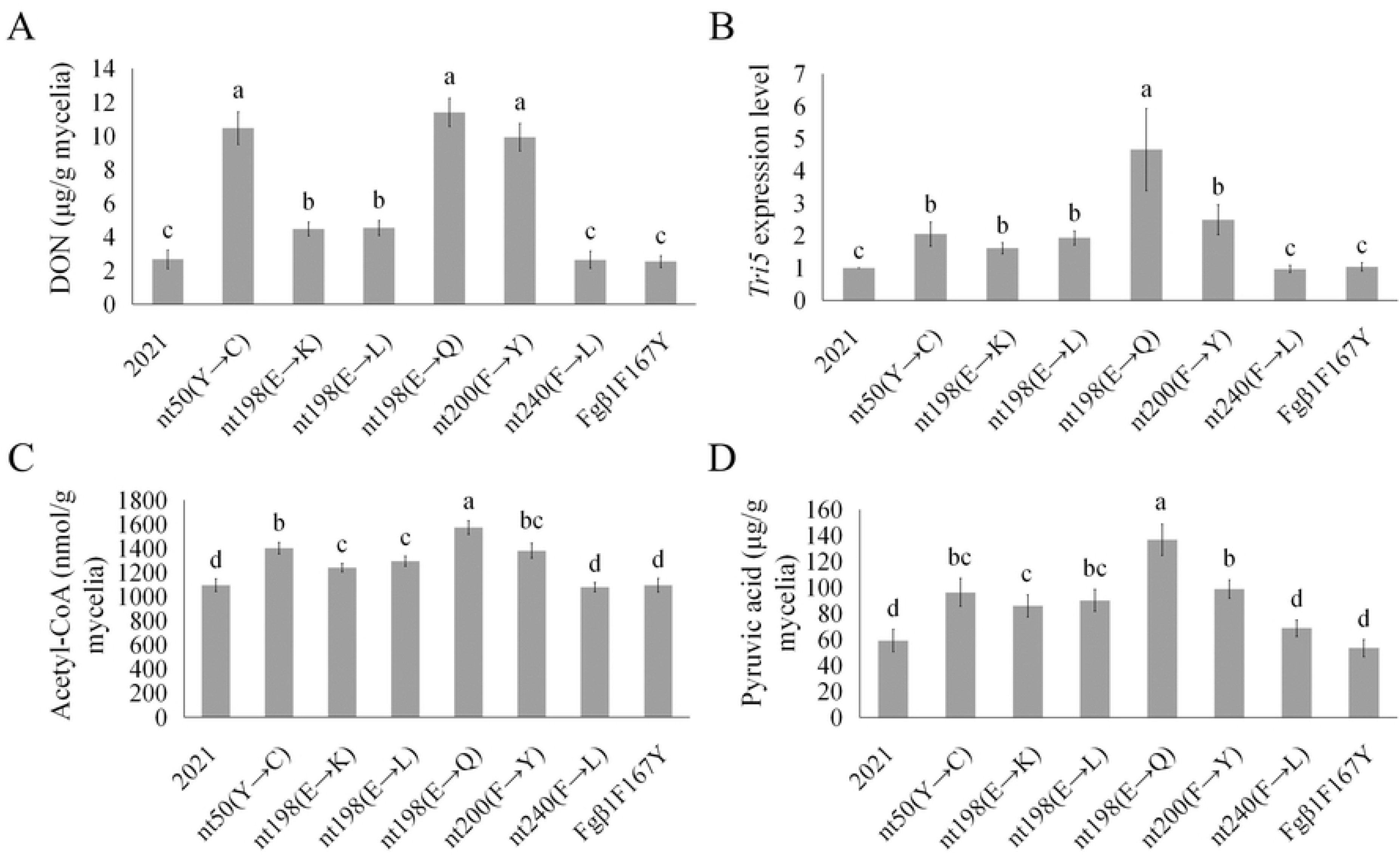
The elevated DON biosynthesis is connected with the carbendazim-resistant substitution of β_2_-tubulin. A, Comparisons of DON production among the wild-type strain 2021 and various β_2_- tubulin mutation mutants. The carbendazim-resistant mutants had increased DON production and *Tri5* expression level while other mutants had no difference in comparison to 2021. Error bar indicates the standard deviation (SD) calculated from the data of three replicates. B, Comparisons of *Tri5* expression levels among the wild-type strain 2021 and various β_2_- tubulin mutation mutants. C, Acetyl-CoA production was increased in carbendazim-resistant substitution mutants in comparison to 2021. Error bar indicates standard deviation (SD) calculated from the data of three replicates. D, Pyruvic acid production was increased in carbendazim-resistant mutants in comparison to 2021.

### Carbendazim-resistant substitutions reduce the interaction between β2-tubulin and FgIDH3

Because β_2_-tubulin is an important skelemin in *F. graminearum* [32,33], it must interact with multiple proteins and regulate their functions. Therefore, we conducted an affinity capture-mass spectrometry (ACMS) assay for β_2_-tubulin to identify novel interacting proteins. In brief, the total protein was isolated from the wild-type strain 2021 and co-purified with magnetic beads which bound anti-β_2_ antibody. After incubation, the bound protein was analyzed by mass spectrometry. The β_2_-tubulin deletion mutant ΔFgβ_2_ was used as a negative control. The results showed that IDH3 interacted with β_2_-tubulin. The interactions of β_2_-tubulin with IDH3 was further confirmed by co-immunoprecipitation assays and bimolecular fluorescence complementation (BiFC) assays (Fig 5A and 5B). In addition, we found that most of the FgIDH3-GFP signals merged well with Fgβ_2_-RFP (Fig 5C and 5D). Furthermore, we observed that the F167Y mutation of β_2_-tubulin reduced the interaction between β_2_-tubulin and IDH3 (Fig 5A).

**Fig 5.**
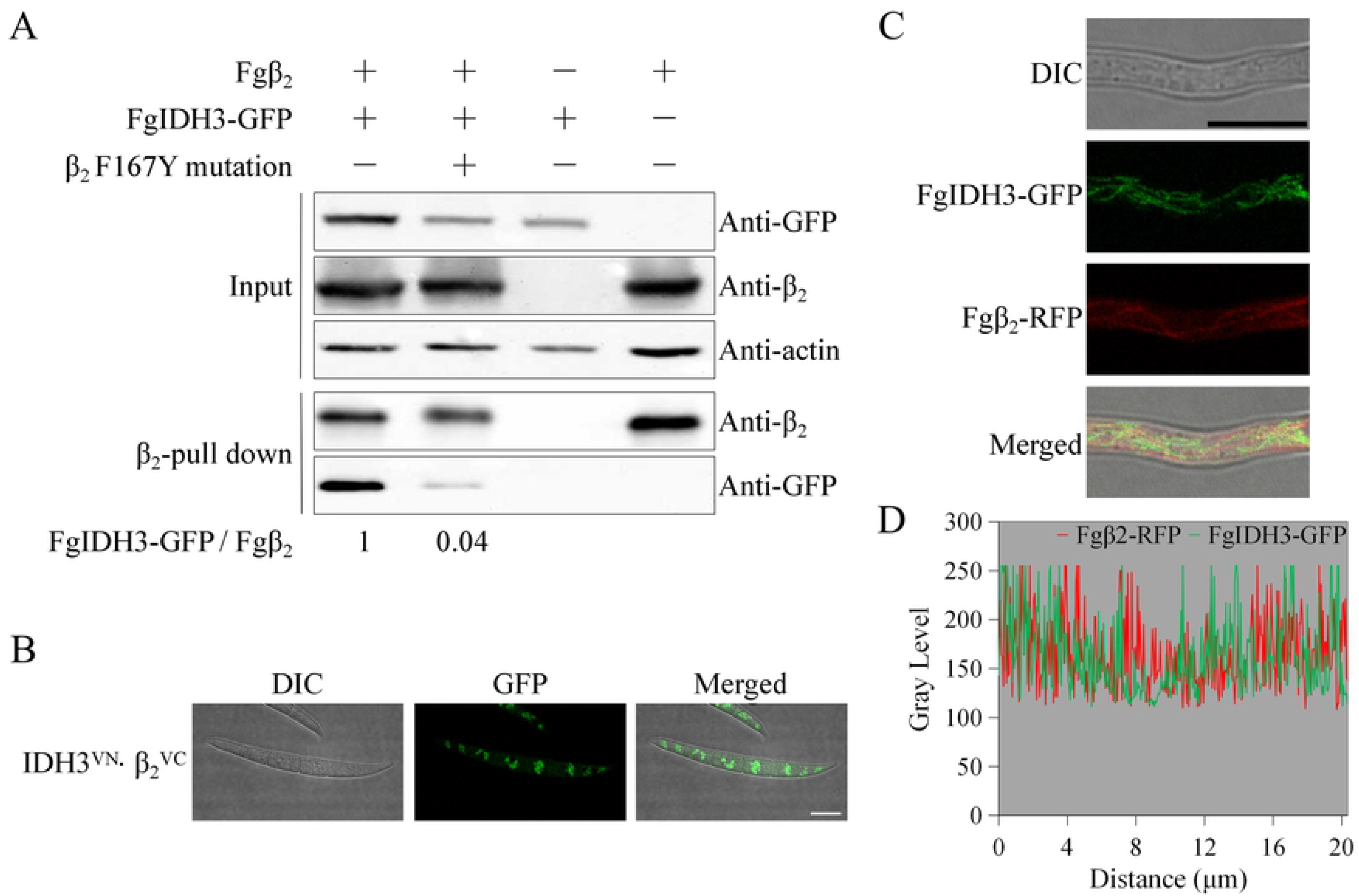
Analysis of the interaction between β_2_-tubulin and IDH3. A, Total proteins (input) extracted from the strain containing Fgβ_2_ and FgIDH3-GFP constructs or a single construct (Fgβ_2_ or FgIDH3-GFP) were subjected to SDS-PAGE and immunoblots were incubated with monoclonal anti-β_2_ and monoclonal anti-GFP antibodies as indicated (upper image). In addition, each protein sample was pulled down using anti-GFP antibodies with magnetic bead and further detected with monoclonal anti-GFP and monoclonal anti-β_2_ antibodies (lower image). The protein samples were also detected with monoclonal anti-β-actin antibody as a reference. B, The interaction of β_2_-tubulin with IDH3 was confirmed by bimolecular fluorescence complementation (BiFC) analysis. The constructs of pFgIDH3-GFPN and pFgβ_2_-GFPC were co-transformed into 2021 to generate the strain FgIDH3-GFPN+Fgβ_2_-GFPC. The GFP signals in the conidia of the strain grown in the MBB medium were examined under a confocal microscope. Bar = 10 μm. C and D, The observation and linescan graph analysis of the co-localization of FgIDH3-GFP and Fgβ_2_-RFP. Bar = 10 μm.

Given that increased DON biosynthesis and acetyl-CoA concentration correlate with carbendazim-resistant substitutions of β_2_-tubulin, we further measured the expression of IDH3 in each carbendazim-resistant mutant. The results showed that the expression of IDH3 in transcriptional and translational levels significantly decreased as compared with the wild-type strain 2021 (Fig 6A and 6B). Since the F167Y mutation of β_2_-tubulin reduces the interaction between β_2_-tubulin and IDH3, a similar phenomenon might exist in other types of carbendazim-resistant mutants. In order to test this hypothesis, we tagged FgIDH3 with GFP in Fgβ_2_^E198Q^, Fgβ_2_^E198L^, Fgβ_2_^E198K^ and Fgβ_2_^F200Y^ mutants to determine the interaction intensity between β_2_-tubulin and IDH3. As expected, the interaction intensity exhibited a significant decrease in the tested carbendazim-resistant mutants (Fig 6C). In addition, we measured the concentrations of citric acid, the upstream substrate of IDH3. Results showed that the production of citric acid significantly increased in the tested carbendazim-resistant mutants (S8A Fig) Moreover, the transcriptional and translational levesls of FgIDH3 significantly decreased in the mutant ΔFgβ_2_ (Fig 5A, S7 Fig). Taken together, these results demonstrated that β_2_-tubulin can increase its expression by interacting with IDH3, and carbendazim-resistant substitutions can reducing the interaction between β_2_-tubulin and IDH3, which resulted in the reduced expression of IDH3.

**Fig 6.**
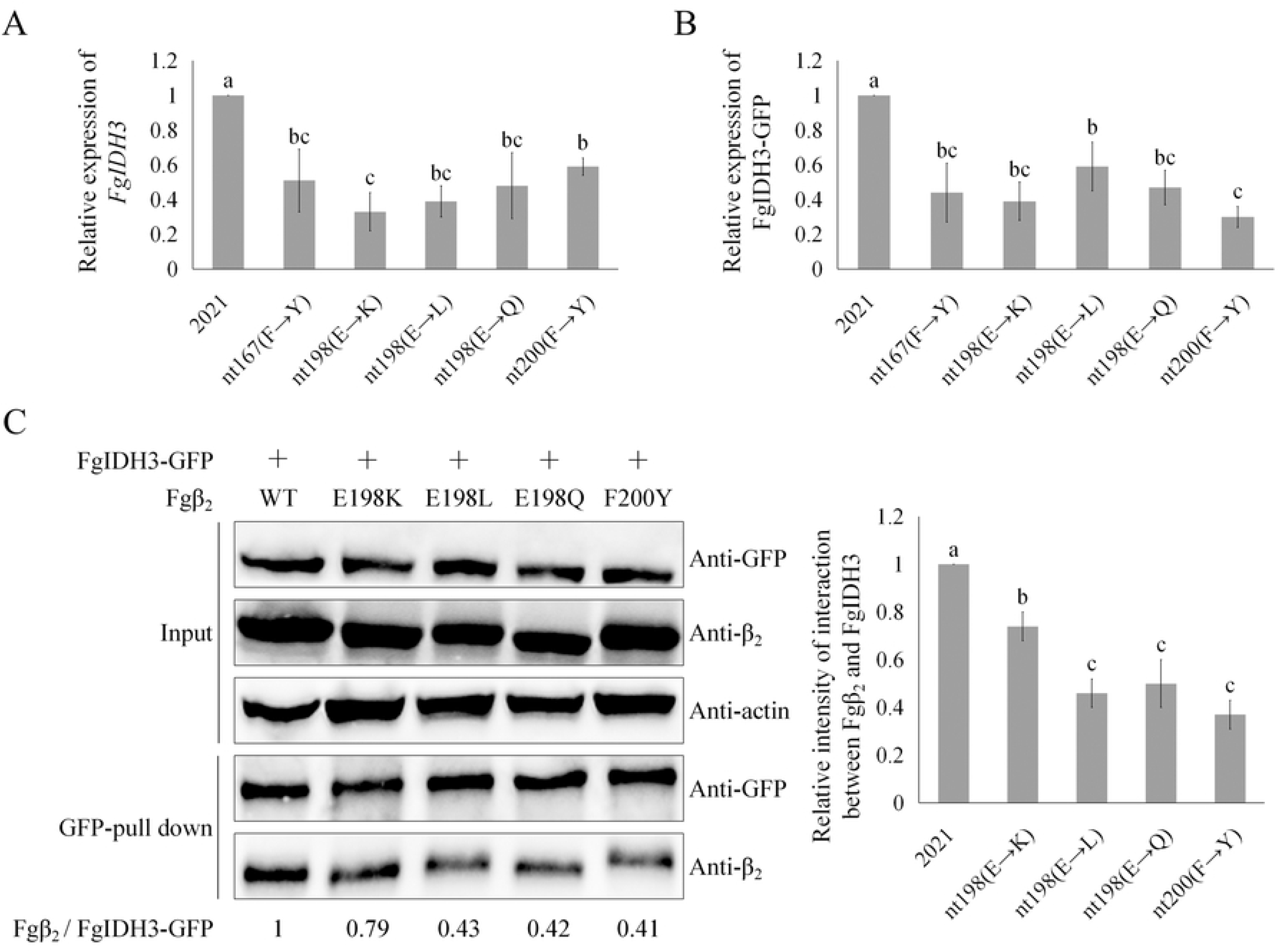
Carbendazim-resistant substitutions reduces FgIDH3 expression by altering the interaction intensity between β_2_-tubulin and IDH3. A and B, The expression of FgIDH3 was detected by qRT-PCR and western blotting assays. Error bar indicates standard deviation (SD) calculated from the data of three replicates. C, The interaction intensity between β_2_-tubulin and IDH3 was tested with the wild-type strain 2021 and carbendazim-resistant mutants were derived. Protein samples were eluted from anti-GFP beads and detected with the anti-GFP and anti-β_2_ antibodies respectively.

### Carbendazim increases DON biosynthesis by disrupting the binding between β_2_-tubulin and IDH3

Because carbendazim-resistant substitutions reduce the interaction intensity between β_2_-tubulin and IDH3, we further examined whether carbendazim has an impact on the interaction of β_2_-tubulin and IDH3. The results showed that the interaction intensity between β_2_-tubulin and IDH3 significantly decreased in the wild-type strain 2021 after being treated with 0.5, 5 and 50 μg/ml (approximately EC_50_, 10×EC_50_ and 100×EC_50_) carbendazim (Fig 7A, S9A Fig). Similarly, all the Fgβ_2_ mutants (Fgβ_2_ ^E167Y^, Fgβ_2_ ^E198K^ and Fgβ_2_ ^F200Y^) also decreased the interaction intensity between β_2_-tubulin and IDH3 after treatment with EC_50_, 10× EC_50_ and 100× EC_50_ carbendazim (Fig 7B-7D, S9B-S9D Fig).

**Fig 7.**
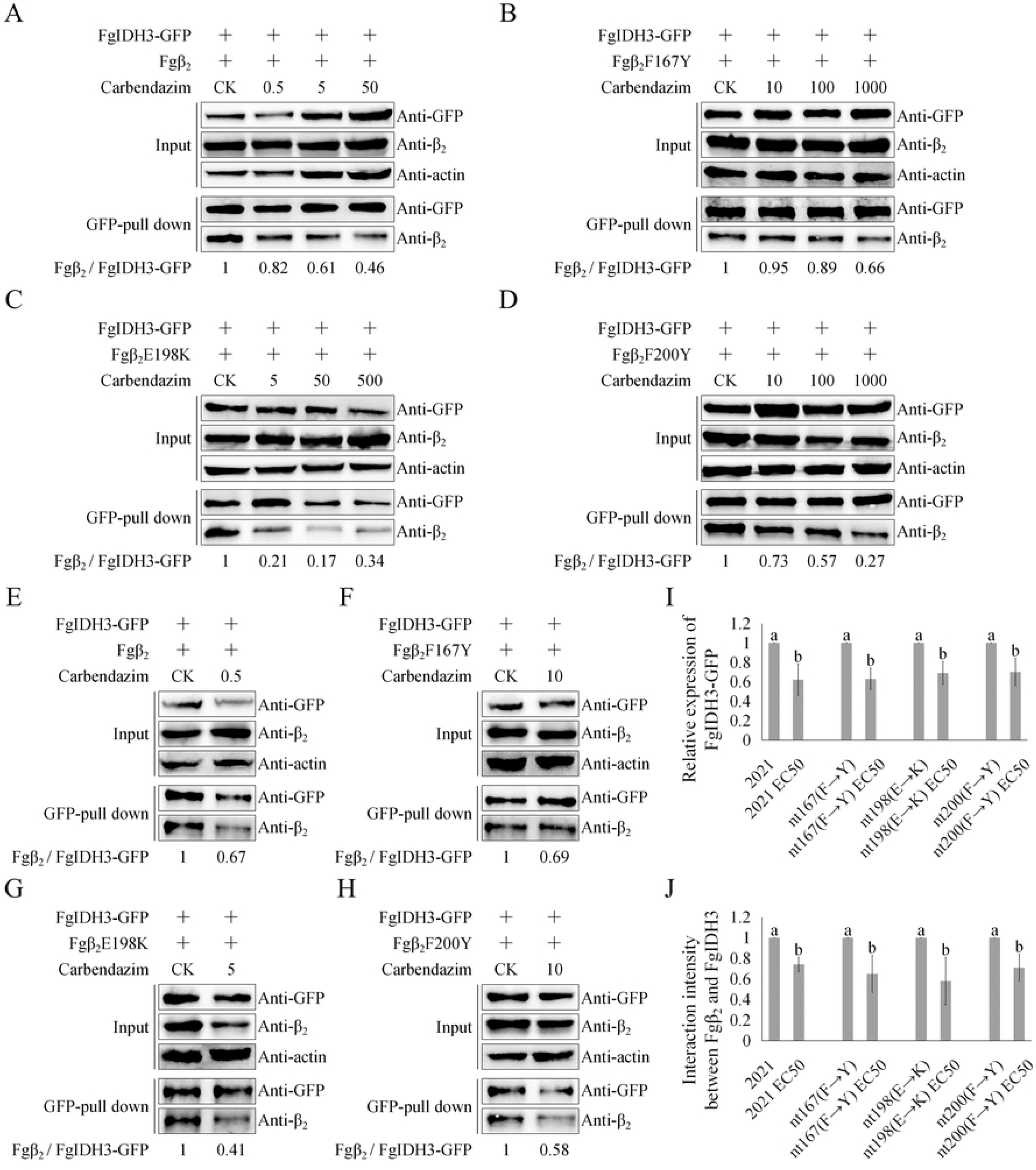
Carbendazim regulates DON biosynthesis by disrupting the interaction between β_2_-tubulin and FgIDH3. A to D, Carbendazim reduced the interaction between β_2_-tubulin and IDH3. Mycelia from the wild-type strain 2021 or the carbendazim-resistant mutants which co-express the Fgβ_2_ and FgIDH3-GFP were treated with carbendazim (EC_50_, 10× EC_50_ and 100× EC_50_) for 3 h and then harvested for protein extraction. Next, the protein samples were eluted from anti-GFP beads and detected with the anti-GFP and anti-β_2_ antibodies respectively. E to H, Carbendazim reduced the interaction between β_2_-tubulin and IDH3 in toxin inducing cultures. I, The expression of FgIDH3-GFP was assayed by western blotting assays with protein extracted from the same set of samples used in 7E to 7H. J, The interaction intensity between β_2_-tubulin and IDH3 was assayed by western blotting assays with protein extracted from the same set of samples used in 7E to 7H. Error bar indicates standard deviation (SD) calculated from the data of three replicates.

Previous studies have shown that carbendazim (approximately EC_50_ against mycelial growth) can accelerate DON biosynthesis in *F. graminearum* [34]. To further understand the role of carbendazim in regulating DON biosynthesis, we treated the wild-type strain 2021 and the β_2_-tubulin mutants (Fgβ_2_^E167Y^, Fgβ_2_^E198K^ and Fgβ_2_^F200Y^) with carbendazim. Briefly, each strain was grown in GYEP medium for 24 h before carbendazim (approximately the EC_50_ of each strain) was added. After incubation for another 48 h, the expression of IDH3 and the interaction intensity between β_2_-tubulin and IDH3 were measured. As indicated, in each strain, the interaction intensity between β_2_-tubulin and IDH3 significantly decreased after treatment with carbendazim in each strain (Fig 7E-7H, 7J). Additionally, western blotting assays showed that the translational level of IDH3 also significantly decreased after treatment with carbendazim (Fig 7I). In addition, the concentrations of citric acid significantly increased after treatment with carbendazim (S8B Fig). Furthermore, we assayed the *Tri5* expression by qRT-PCR. As expected, *Tri5* was up-regulated after being treated with carbendazim in the wild-type strain 2021 and the β_2_-tubulin mutants (data not shown). These results indicated that carbendazim can reduce the interaction between β_2_-tubulin and IDH3, decreasing the expression of IDH3 and thus accelerating DON biosynthesis in *F. graminearum*.

## Discussion

Fusarium head blight (FHB) is one of the most destructive diseases of wheat worldwide, and the mycotoxins produced by FHB pathogens are carcinogenic to humans and animals [35,36]. Despite being the major way of controlling FHB, the fungicides currently used cannot effectively control DON contamination. Moreover, some fungicides such as carbendazim and their improper usage in the field not only give rise to fungicide resistance but also trigger certain cellular metabolism changes in fungi, such as the stimulation of DON biosynthesis [4,22,34]. Recently, the mutation at codon 167 of β_2_-tubulin was detected as the most frequent type in the field and can significantly increase DON biosynthesis in *F. graminearum* [22]. Transcriptome assays revealed that the mutant nt167 had global gene expression difference as compared to the wild-type strain 2021, including acetyl-CoA metabolism. In our current studies, we demonstrated that the expression of the regulatory enzymes in the TCA cycle was down-regulated, such as FgIDH1, FgIDH2 and FgIDH3 (Fig 1A). In addition, we found that intracellular acetyl-CoA significantly increased in nt167. In order to understand the molecular mechanism behind the increased DON biosynthesis in *F. graminearum*, we conducted an affinity capture-mass spectrometry (ACMS) assay for β_2_-tubulin, so as to identify novel interaction proteins. Interestingly, the masses of enzymes involved in acetyl-CoA metabolism were identified as Fgβ_2_-interacting proteins, including aldehyde dehydrogenase, acetyl-CoA synthetase, pyruvate decarboxylase and isocitrate dehydrogenase. These preliminary data suggest that the carbendazim-resistant mutation of β_2_-tubulin, might regulate DON biosynthesis by altering the expression of metabolic enzymes involved in acetyl-CoA metabolism in *F. graminearum*.

As a central metabolite and second messenger, acetyl-CoA plays a vital role in the regulation of gene expression related to histone acetylation, cell growth and mitosis, catabolism and autophagy, caspase-dependent apoptosis and regulated necrosis [37–41]. In this study, we revealed that DON biosynthesis was induced by exogenous acetyl-CoA. The results showed that DON production and *TRI*5 expression in the wild-type 2021 significantly increased after treatment with 8 μg/ml acetyl-CoA (Fig 1D and 1E). Although intracellular acetyl-CoA has been implicated in the regulation of secondary metabolism in the industrial microorganism *Saccharomyces cerevisiae* [8–10], to our knowledge significant stimulation of DON biosynthesis by exogenous acetyl-CoA has not been reported in plant pathogenic fungi. However, exogenous acetyl-CoA barely had any effect on DON biosynthesis before *TRI* genes transcription, even after treatment with 20 μg/ml acetyl-CoA. It is possible that exogenous acetyl-CoA was used for other metabolic pathways related to vegetative growth and cell proliferation before *TRI* genes transcription. In addition, we observed that exogenous acetyl-CoA stimulated the formation of bulbous hyphae associated with DON biosynthesis after *TRI* genes transcription (Fig 1F). In eukaryotic cells, acetyl-CoA is known to regulate cell growth and proliferation [42]. In budding yeast, elevated acetyl-CoA concentration could induce cell growth by promoting the acetylation of histones which are required for growth [36,43]. Embryonic stem cells rapidly lose pluripotency accompanied by a metabolic shift including the decreased glycolytic activity, lower intracellular acetyl-CoA availability as well as histone deacetylation [41]. The exogenous supply of acetate, the substrate of acetyl-CoA synthetase, could efficiently delay embryonic stem cell differentiation [41]. It is possible that the cellular differentiations related to DON biosynthesis regulated by acetyl-CoA in *F. graminearum* might be associated with the acetylation of histones connected with *TRI* genes expression.

In eukaryotic cells, acetyl-CoA is predominantly synthesized in the mitochondrial matrix and then metabolized within the TCA cycle [42,44]. Our studies showed that FgIDH2 and FgIDH3 are localized in the mitochondria while FgIDH1 is localized in the vacuoles (Fig 2B), indicating that FgIDH2 and FgIDH3 might have a similar biological function. In pancreatic beta cells, the knockdown of mitochondrial IDH resulted in various changes in metabolite levels, including the production of α-ketoglutarate, malate and citrate as well as the release of insulin [11]. Similarly, a previous study with the IDH mutant of *Aspergillus niger* with overexpression of the IDH gene *in vivo* resulted in the reduction of citric acid production [16]. In addition, IDH is required for lipid biosynthesis in melanoma cells under hypoxic conditions [45]. Furthermore, the deficiency of mitochondria IDH activity resulted in mitochondrial dysfunction and cardiac hypertrophy in mice, which were associated with disruption of redox homeostasis [15]. Therefore, the IDH gene is an important regulator, not only in acetyl-CoA metabolism but redox homeostasis of eukaryotic organisms. In our study, the deletion mutants of each isocitrate dehydrogenase gene, IDH2 and IDH3 significantly elevated the intracellular acetyl-CoA and DON biosynthesis in *F. graminearum*. These results indicated that, although IDH1 regulates DON biosynthesis, IDH2 and IDH3 are the major regulators in negatively controlling DON biosynthesis in *F. graminearum*.

In previous studies, MTs were properly characterized in terms of cell polarity, mitosis, motility and vesicle traffic. In addition, the indispensable role of MTs in regulating various metabolic processes have been reported [23]. Almost all glycolytic enzymes can bind to MTs to a certain degree. For example, enolase, one of the glycolytic enzymes, interacts with MTs during various stages of cell differentiation [46]. A previous study reported that MTs could bind glyceraldehyde 3-phosphate dehydrogenase (GAPDH) and then regulate its activity and quaternary structure [47]. In addition to directly modulating the three-dimensional structure of enzymes, MTs have been thought to facilitate enzymatic catalytic reactions by serving as a scaffold for organizing different enzymes in sequential order. In HeLa cells, MTs network is indispensable for the spatial distribution/assembly of purinosomes, which is beneficial to purine synthesis [48]. MTs can accelerate the kinase activity of Aurora-B by reducing its dimensionality [49]. Furthermore, MTs can act as the transcription factor of some proteins. Under hypoxic conditions, MTs are the prerequisite for activating the translation of hypoxia inducible factor (HIF)-1ɑ, which provide the attachment points of polysomes associated with HIF-1ɑ mRNA translation [50]. In our study, we demonstrated that β_2_-tubulin physically binds with the acetyl-CoA-associated enzyme IDH3, like various glycolytic enzymes mentioned above, thereby increasing the expression of IDH3 in the mitochondria and further participating in the regulation of DON biosynthesis in *F. graminearum*. Although several researches have shown that tubulin dimers regulate the physiological function of the mitochondia by binding the voltage-dependent anion channel (VDAC), one of the mitochondrial outer membrane proteins, no evidences indicated that tubulin exists in the mitochondrial matrix and inner membrane [23]. Consequently, the interaction between β_2_-tubulin and IDH3 most likely happened before the newly synthesized IDH3 proteins were transported into the mitochondria and the transportation process is known as post-translational translocation. It is possible that β_2_-tubulin plays an important role in assisting IDH3 spatial structure formation, similar to the molecular chaperones. Thus, it will be interesting to study the interaction mode between β_2_-tubulin and IDH3 and reveal a new function of β_2_-tubulin in regulating carbon metabolism.

Apart from the biological functions of MTs, the point mutation in MTs has also been extensively studied in eukaryotic organisms. Evidences have shown that human *TUBB3* disease-associated mutations can impair tubulin heterodimer formation, disrupt the interaction of tubulin with kinesin motors and result in defection in axon guidance [32]. Two disease-associated mutations in *TUBB2B* can result in asymmetrical polymicrogyria by disrupting microtubule-based processes [51]. In *S. cerevisiae*, mutation in *TUB2* can inhibit the mitotic cell cycle and block chromosome segregation [52]. Furthermore, a point mutation in *TUBB3* which is close to the kinesin-binding site, can alter the polymerization dynamics of MTs [53]. In the current study, we found that the carbendazim-resistant mutations (F167Y, E198K, E198L, E198Q and F200Y) of β_2_-tubulin can reduce the interaction intensity between β_2_-tubulin and the acetyl-CoA-associated enzyme IDH3, and decrease the expression of IDH3 in the mitochondria, resulting in the accumulation of cytosolic acetyl-CoA as well as the increase of DON biosynthesis in *F. graminearum*. Besides, our findings revealed that carbendazim can also disrupt the interaction between β_2_-tubulin and IDH3. These results indicated that the IDH3 binding domain on β_2_-tubulin might be associated with carbendazim-resistant mutation sites. Furthermore, we demonstrated that the stimulation of DON biosynthesis affected by carbendazim is also caused by reduction of decreased interaction intensity between β_2_-tubulin and IDH3. Thus, the selection of fungicides for FHB control, especially carbendazim and other benzimidazoles, must be decided after deliberate consideration, to avoid not only widespread drug resistance but the stimulation of mycotoxin biosynthesis. Taken together, our data support a model that carbendazim-resistant substitutions of β_2_-tubulin can increase DON biosynthesis by reducing the interaction between β_2_-tubulin and IDH3, and carbendazim can cause the stimulation of DON by disrupting the binding between β_2_-tubulin and IDH3 in *F. graminearum* (Fig 8).

**Fig 8.**
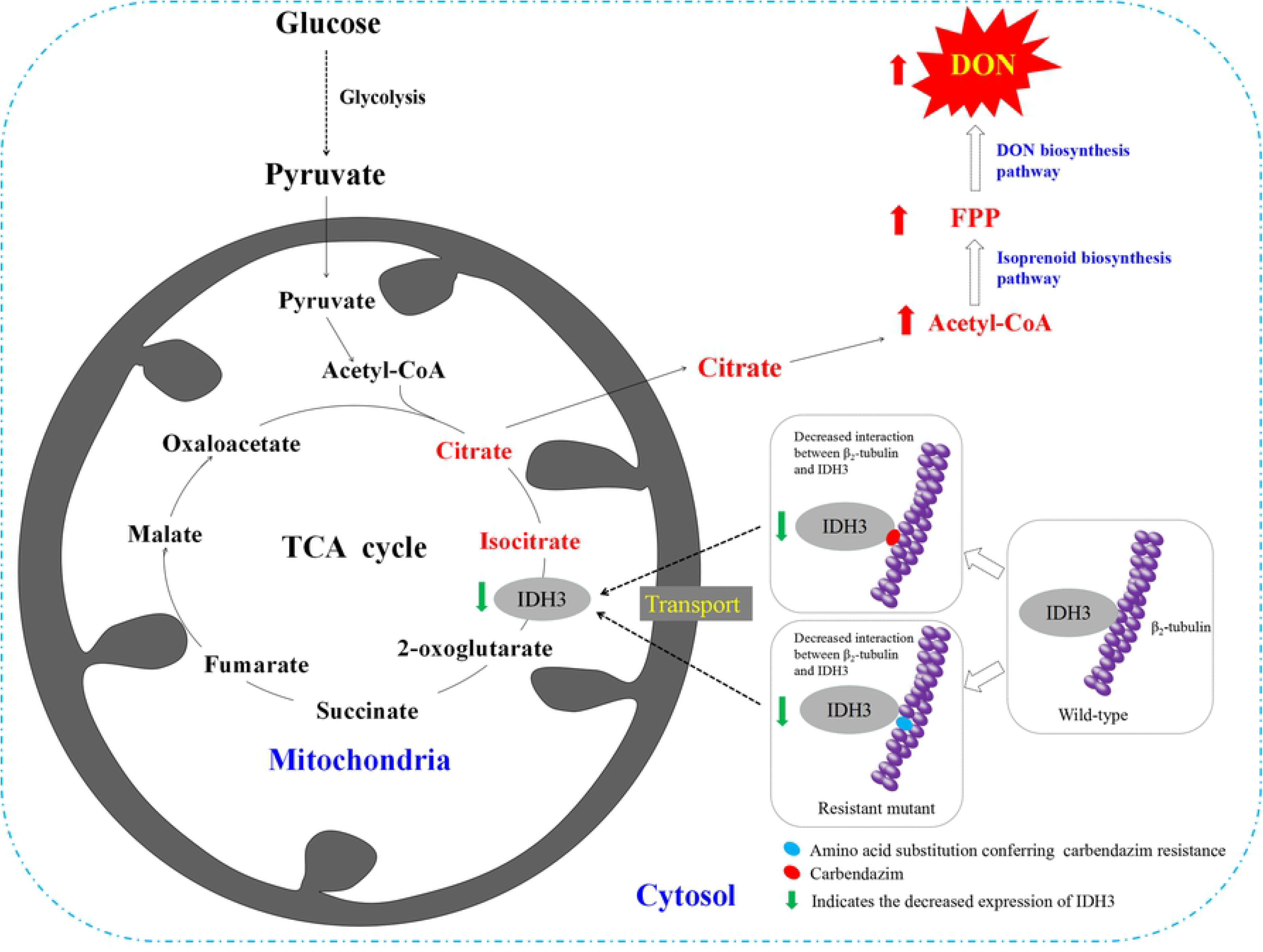
Proposed model for carbendazim-resistant substitutions regulating DON biosynthesis in *F. graminearum*. β_2_-tubulin can bind isocitrate dehydrogenase subunit 2 (IDH3) and regulates its expression in *F. graminearum*. In carbendazim-resistant strains, carbendazim-resistant substitutions reduced the interaction intensity between β_2_-tubulin and IDH3. In addition, β_2_-tubulin inhibitor carbendazim is able to disrupt the binding between β_2_-tubulin and IDH3. Furthermore, the decreased interaction intensity between the β_2_-tubulin and IDH3 caused the reduced expression of IDH3, thereby stimulating the biosynthesis of DON in *F. graminearum*.

## Materials and Methods

### Strains and culture conditions

The *F. graminearum* carbendazim-sensitive strain 2021 was used as a wild-type strain. Growth rate and colony morphology assays of the strains generated in this research were grown on potato dextrose agar (PDA). The Mung bean broth (MBB) was used for conidia culture as described previously [54]. For DON production analysis and bulbous structures observation, all strains were grown in GYEP medium at 28°C in dark.

### Strain construction

All gene deletion mutants were constructed using the protocols described previously [54]. The open reading frame (ORF) of each gene was replaced with *HPH-HSV-tk*, the transformants were identified by PCR assays and confirmed by Southern blot analysis.

To construct FgIDH3-GFP cassette, FgIDH3 containing the native promoter region and ORF (excluding the stop codon) was amplified. Then the PCR product was co-transformed with *XhoI*-digested pYF11 into the yeast strain XK1-25 as described [55]. Subsequently, the FgIDH3-GFP fusion vector was recovered from the yeast transformant by using the Yeast Plasmid Kit (Solarbio, Beijing, China) and then transferred into *E. coli* strain DH5α for amplification. With the similar strategy, other GFP fusion cassettes were also constructed. Each plasmid was transformed into the wild-type strain 2021.

### Western blotting assays

Each tested strain was incubated in YEPD at 25°C for 2 days, then mycelia was harvested and washed with sterile water for protein extraction. About 100 mg of liquid nitrogen grinded mycelia was re-suspended in 1 ml of extraction buffer [50 mM Tris–HCl, pH 7.5, 100 mM NaCl, 5 mM ethylene diamine tetraacetic acid (EDTA), 1% Triton X-100, 2 mM phenylmethylsulfonyl fluoride (PMSF)] and 10 μl of protease inhibitor cocktail (Sangon, Shanghai, China). After homogenization with a vortex shaker, the lysate was centrifuged at 12000 g in a microcentrifuge for 20 min at 4°C. 15μl of each sample was loaded onto 10% SDS-PAGE gels. Proteins separated on gels were transferred to Immobilon-P transfer membrane (Millipore, Billerica, MA, USA). The monoclonal anti-GFP antibody 300943 (Zenbio, Sichuan, China) and polyclonal anti-β_2_ antibody (IF11) [56] were used at a 1:1000 and 1:20000 dilution for immunoblot analyses, respectively. The samples were also detected with monoclonal anti-β-actin antibody 700068 (Zenbio, Sichuan, China) as a reference. Incubation with a secondary antibody and chemiluminescent detection were performed as described previously [57].

### Affinity capture-mass spectrometry analysis

The wild-type strain 2021 was used for protein extraction as described previously [58]. After protein extraction, supernatant was transferred into a sterilized tube. The affinity capture was conducted as described. First, 50 μl magnetize beads (Bio-Rad Co., California, USA) were incubated with 0.6 μl IF11 in final volume of 300 μl PBS-T [PBS (137 mM NaCl, 2.7 mM KCl, 10 mM Na_2_HPO_4_, 2 mM KH_2_PO_4_, PH 7.4) + 0.1% Tween 20] in a sterilized eppendorf tube and rotate 10 minutes at room temperature. Second, magnetized beads and discard supernatant, then washed beads with 1 ml PBS-T for three times. Third, transferred protein supernatant (1mL) into the tube and rotated for 1 hour at room temperature to capture β_2_-tubulin interacting proteins. The last step is discard supernatant and washed beads with 1 ml PBS-T for three times. To identify β_2_-tubulin interacting proteins, the samples were washed and further analyzed by MS analysis (Shanghai Applied Protein Technology Co., Ltd.).

### Co-immunoprecipitation (Co-IP) assays

The GFP-fusion constructs was verified by DNA sequencing and transformed into 2021. Transformants expressing β_2_-tubulin and GFP-fusion protein were confirmed by western blotting analyses. Besides, the transformants expressing β_2_-tubulin or GFP-fusion protein were used as references. For Co-IP assays, total protein was extracted and incubated with the anti-GFP antibody as described above. Protein eluted from magnetic bead was analysed by western blot detection with IF11 or anti-GFP antibody. The protein sample was also detected with anti-β-actin antibody as a reference.

### Bimolecular Fluorescence Complementation (BiFC) assays

For BiFC assays, the plasmid constructs of pGFPN-FgIDH3 and pFgβ_2_-GFPC were verified by sequencing and then co-transformed into carbendazim-sensitive strain 2021 in pairs. Transformants resistant to both neomycin and hygromycin were isolated and confirmed by PCR. The recombination plasmid pGFPN-FgIDH3 or pFgβ_2_-GFPC was transformed into 2021, and the correct transformant was used as negative controls. GFP signals in the conidia grown in MBB medium 3 day were examined under a Leica TCS SP5 confocal microscope (Wetzlar, Hessen, Germany).

### Assays for stimulation effects of acetyl-CoA on DON biosynthesis and TRI gene expression

Acetyl-CoA (Solarbio, Beijing, China) was added to a final concentration of 1, 2, 4 or 8 μg/ml to GYEP culture after a 3-day incubation. DON production in GYEP cultures was assayed with a competitive ELISA based DON detection plate kit (Wise, Zhenjiang, China) according to the previous studies [59,60].

To assaying *TRI* gene expression, hyphae was harvested from 4-day-old GYEP cultures (1 day after acetyl-CoA were added) and used for RNA isolation with the RNAsimple Total RNA Kit (Tiangen, Beijing, China), and cDNA was synthesis by HiScript II Q RT SuperMix for qPCR kit (Vazyme Biotech, Nanjing, China) following the protocols and used for qPCR with ChamQ^™^ SYBR qPCR Master Mix kit (Vazyme Biotech, Nanjing, China) following manufacturer’s instructions. The actin gene of *F. graminearum* was used as the internal control. The relative quantification of each gene was calculated with the 2^−ΔΔCt^ method [61]. The results were calculated with the data from three biological replicates.

### Microscopic examinations

Hyphal morphology was examined with an Olympus IX-71 microscope (Tokyo, Japan) using fresh mycelia taken from 1-day-old colonies grown on PDA plates. The localization of tagged protein was observed with a Leica TCS SP5 confocal microscope (Wetzlar, Hessen, Germany). Each strain was cultured in YEPD for 1 day before staining with different fluorescent Dyes. The following filter sets were used for fluorescent or dye staining: the laser excitation wavelength was set at 488 nm for GFP-fusion proteins (green fluorescence), at 353 nm for CMAC (blue fluorescence). The mitochondria was stained with Mito Tracker Red CMXRos (Yeasen Co., Ltd), and laser was set at 579 nm for red fluorescence.

### Quantification of acetyl-CoA and citric acid production

The mycelia grown in YEPD for 2 days and in GYEP for 3 days were used for the measurement of acetyl-CoA and citric acid. Acetyl-CoA and citric acid production was assayed using an Acetyl-CoA Assay Kit (Solarbio, BC0980) and a Citric acid Assay Kit (Solarbio, BC2155), respectively. Briefly, mycelia (0.05 g) were added to 500 mL of lysis buffer in the acetyl-CoA or citric acid detection kit. After lysis of mycelia, quantification of acetyl-CoA and citric acid production was conducted following the protocols provided by the manufacturer. The experiments were repeated three times.

## Acknowledgements

We appreciate Prof. WB Shim at Texas A&M University on the manuscript editing. We thank Prof. JR Xu at Purdue University for providing the PYF11 GFP-plasmid used in this study. This work was supported by the Key Program of National Natural Science Foundation of China 31730072 (to M.Z.) and the General Program of National Natural Science Foundation of China 31772190 (to Y.D.).

## Supporting information

**S1 Fig. Effects of exogenous acetyl-CoA on DON biosynthesis in the ΔFghxk mutant.** A, DON production in GYEP cultures of ΔFghxk treatment with 0, 1, 2 and 4 μg/ml acetyl-CoA after incubation for 7 days. Error bar indicates standard deviation (SD) calculated from the data of three replicates. B, *Tri5* expression assayed by qRT-PCR with RNA sample isolated from ΔFghxk which cultures incubated in the presence/absence of 4 μg/ml acetyl-CoA for 4 days. Acetyl-CoA was added after a 3-day incubation. The relative expression level of *Tri5* in cultures without exogenous acetyl-CoA was set to 1. Error bar indicates standard deviation (SD) calculated from the data of three replicates. C, GYEP cultures of the ΔFghxk mutant treatment with or without 4 μg/ml acetyl-CoA were examined for bulbous structures (marked with arrows) after acetyl-CoA added for 1, 2 and 3 days. Bar = 20 μm.

**S2 Fig. Effects of exogenous acetyl-CoA with different treatment points on DON production in ΔFghxk and 2021.** ΔFghxk and 2021 treatment with 4 and 8 μg/ml acetyl-CoA in 0, 1, 2 and 3 day, respectively, DON production were assayed after incubation for 7 days. Bars denote standard errors from three repeated experiments. Values on the bars followed by the same letter are not significantly different at P = 0.05 according to a Fisher’s LSD test.

**S3-S5 Figs. Targeted gene replacement of FgIDHs in *F***. *graminearum*.

**S6 Fig. Mycelial morphology of the wild-type strain 2021 and ΔFgIDHs mutants in *F***. *graminearum*.

**S7 Fig. The expression level of FgIDH3 in the wild-type strain 2021 and ΔFgβ_2_ mutant in *F*. *graminearum***.

**S8 Fig Carbendazim-resistant mutation and carbendazim increase citric acid production.** A, Carbendazim-resistant mutation increase citric acid production. B, Carbendazim treatment increase citric acid production in 2021 and derived carbendazim-resistant mutants.

**S9 Fig. The interaction intensity between β_2_-tubulin and IDH3 in the wild-type strain 2021 and derived carbendazim-resistant mutants after treatment with carbendazim**.

**S1 Table. Wild-type and mutant strains of *F. graminearum* used in this study**.

**S2 Table. Primers used in this study**.

**S3 Table. Defects of mutants in growth, conidiation, carbendazim sensibility and DON production**.

## References

1. Goswami RS, Kistler HC. 2004. Heading for disaster: *Fusarium graminearum* on cereal crops. Mol Plant Pathol. 2004; 5: 515–525.

2. Alexander NJ, Proctor RH, McCormick SP. Genes, gene clusters, and biosynthesis of trichothecenes and fumonisins in *Fusarium*. Toxin Reviews. 2009; 28: 198–215.

3. De Walle JV, Sergent T, Piront N, Toussaint O, Schneider YJ, Larondelle Y. Deoxynivalenol affects in vitro intestinal epithelial cell barrier integrity through inhibition of protein synthesis. Toxicol Appl Pharmacol. 2010; 245: 291–298.

4. Audenaert K, Callewaert E, Höfte M, De Saeger S, Haesaert G. Hydrogen peroxide induced by the fungicide prothioconazole triggers deoxynivalenol (DON) production by *Fusarium graminearum*. BMC Microbiol. 2010; 10: 112.

5. Brown DW, Dyer RB, McCormick SP, Kendra DF, Plattner RD. Functional demarcation of the *Fusarium* core trichothecene gene cluster. Fungal Genet Biol. 2004; 41: 454–462.

6. Kimura M, Tokai T, Takahashi-Ando N, Ohsato S, Fujimura M. Molecular and genetic studies of *Fusarium* trichothecene biosynthesis: pathways, genes, and evolution. Biosci Biotechnol Biochem. 2007; 71: 2105–2123.

7. Nitschke W, Russell MJ. Beating the acetyl coenzyme A-pathway to the origin of life. Philos Trans R Soc Lond B Biol Sci. 2013; 368: 20120258.

8. Shiba Y, Paradise EM, Kirby J, Ro DK, Keasling DJ. Engineering of the pyruvate dehydrogenase bypass in *Saccharomyces cerevisiae* for high-level production of isoprenoids. Metab Eng. 2007; 9: 160–168.

9. Chen Y, Daviet L, Schalk M, Siewers V, Nielsen J. Establishing a platform cell factory through engineering of yeast acetyl-CoA metabolism. Metab Eng. 2013; 5: 48–54.

10. Krivoruchko A, Serrano-Amatriain C, Chen Y, Siewers V, Nielsen J. Improving biobutanol production in engineered *Saccharomyces cerevisiae* by manipulation of acetyl-CoA metabolism. J Ind Microbiol. 2013; 40: 1051–1056.

11. MacDonald MJ, Brown LJ, Longacre MJ, Stoker SW, Kendrick MA, Hasan NM. Knockdown of both mitochondrial isocitrate dehydrogenase enzymes in pancreatic beta cells inhibits insulin secretion. Biochim Biophys Acta. 2013; 11: 5104–5111.

12. Israelsen WJ, Heiden MGV. Pyruvate kinase: function, regulation and role in cancer. Semin Cell Dev Biol. 2015; 43: 43–51.

13. Yang ES, Lee SM, Park JW. Silencing of cytosolic NADP^+^-dependent isocitrate dehydrogenase gene enhances ethanol-induced toxicity in HepG2 cells. Arch Pharm Res. 2010; 33: 1065–1071.

14. Kim SH, Yoo YH, Lee JH, Park JW. Mitochondrial NADP+-dependent isocitrate dehydrogenase knockdown inhibits tumorigenicity of melanoma cells. Biochem Biophys Res Commun. 2014; 451: 246–251.

15. Ku HJ, Ahn Y, Lee JH, Park KM, Park JW. IDH2 deficiency promotes mitochondrial dysfunction and cardiac hypertrophy in mice. Free Radic Biol Med. 2015; 80: 84–92.

16. Kobayashi K, Hattori T, Hayashi R, Kirimura K. Overexpression of the NADP^+^-specific isocitrate dehydrogenase gene (icdA) in citric acid-producing *Aspergillus niger* WU-2223L. Biosci Biotechnol Biochem. 2014; 78: 1246–1253.

17. Zhou MG, Ye ZY, Liu JF. Progress of fungicide resistance. J Nanjing Agric Univ. 1994; 17: 33–41.

18. Zhou MG, Wang JX. Study on sensitivity base-line of *Fusarium graminearum* to carbendazim and biological characters of MBC-resistant strains. Acta Phytophysiol Sinica. 2001; 31: 365–370.

19. Chen CJ, Bi CW, Yu JJ, Wang JX, Li HX, Luo QQ, et al. Carbendazim-resistance and its molecular mechanism in *Gibberella zeae*. J Plant Pathol. 2008; 90: 69.

20. Duan YB, Yang Y, Wang JX, Liu CC, He LL, Zhou MG. Development and application of loop-mediated isothermal amplification for detecting the highly benzimidazole-resistant isolates in *Sclerotinia sclerotiorum*. Sci Rep. 2015; 5: 17278.

21. Yang Y, Li MX, Duan YB, Li T, Shi YY, Zhao DL, et al. A new point mutation in β_2_-tubulin confers resistance to carbendazim in *Fusarium asiaticum*. Pestic Biochem Physiol. 2018; 145: 15–21.

22. Zhang YJ, Yu JJ, Zhang YN, Zhang X, Cheng CJ, Wang JX, et al. Effect of carbendazim resistance on trichothecene production and aggressiveness of *Fusarium graminearum*. Mol Plant Microbe Interact. 2009; 22: 1143–1150.

23. Cassimeris L, Silva VC, Miller E, Ton Q, Molnar C, Fong J. Fueled by microtubules: does tubulin dimer/polymer partitioning regulate intracellular metabolism? Cytoskeleton. 2012; 69: 133–143.

24. Walsh J, Keith T, Knull H. Glycolytic enzyme interactions with tubulin and microtubules. Biochim Biophys Acta. 1989; 999: 64–70.

25. Vertessy B, Orosz F, Kovacs J, Ovadi J. Alternative binding of two sequential glycolytic enzymes to microtubules. J Biol Chem. 1997; 272: 25542–25546.

26. Orosz F, Wagner G, Liliom K, Kovacs J, Baroti K, Horanyi M, et al. Enhanced association of mutant triosephosphate isomerase to red cell membranes and to brain microtubules. Proc Natl Acad Sci U S A. 2000; 97: 1026–1031.

27. Wagner G, Kovacs J, Low P, Orosz F, Ovadi J. Tubulin and microtubule are potential targets for brain hexokinase binding. FEBS Lett. 2001; 509: 81–84.

28. Jonkers W, Dong Y, Broz K, Kistler HC. The Wor1-like protein Fgp1 regulates pathogenicity, toxin synthesis and reproduction in the phytopathogenic fungus *Fusarium graminearum*. PLoS Pathog. 2012; 8: e1002724.

29. Menke J, Weber J, Broz K, Kistler HC. Cellular development associated with induced mycotoxin synthesis in the filamentous fungus *Fusarium graminearum*. PLoS One. 2013; 8: e63077.

30. Sakamoto N, Tsuyuki R, Yoshinari T, Usuma J, Furukawa T, Nagasawa H, et al. Correlation of ATP citrate lyase and acetyl CoA levels with trichothecene production in *Fusarium graminearum*. Toxins. 2013; 5: 2258–2269.

31. Ohneda M, Arioka M, Nakajima H, Kitamoto K. Visualization of vacuoles in *Aspergillus oryzae* by expression of CPY-EGFP. Fungal Genet Biol. 2002; 37: 29–38.

32. Tischfield MA, Baris HN, Wu C, Rudolph G, Maldergem LV, He W, et al. Human *TUBB3* mutations perturb microtubule dynamics, kinesin interactions, and axon guidance. Cell. 2010; 140: 74–87.

33. Chuong S, Good A, Taylor G, Freeman M, Moorhead G, Muench D. Large-scale identification of tubulin-binding proteins provides insight on subcellular trafficking, metabolic channeling, and signaling in plant cells. Mol Cell Proteomics. 2004; 3: 970–983.

34. Tang GF, Chen Y, Xu JR, Kistler HC, Ma ZH. The fungal myosin I is essential for *Fusarium* toxisome formation. PLOS Pathog. 2018; 14: e1006827.

35. Placinta C, D’mello J, Macdonald A. A review of worldwide contamination of cereal grains and animal feed with *Fusarium* mycotoxins. Anim Feed Sci Technol. 1999; 78: 21–37.

36. Cleveland TE, Dowd PF, Desjardins AE, Bhatnagar D, Cotty PJ. United States Department of Agriculture-Agricultural Research Service research on pre-harvest prevention of mycotoxins and mycotoxigenic fungi in US crops. Pest Manag Sci. 2003; 59: 629–642.

37. Wellen KE, Hatzivassiliou G, Sachdeva UM, Bui TV, Cross JR, Thompson CB. ATP-citrate lyase links cellular metabolism to histone acetylation. Science. 2009; 324: 1076–1080.

38. Cai L, Sutter BM, Li B, Tu BP. Acetyl-CoA induces cell growth and proliferation by promoting the acetylation of histones at growth genes. Mol Cell. 2011; 42: 426–437.

39. Eisenberg T, Schroeder S, Andryushkova A, Pendl T, Kuttner V, Bhukel A, et al. Nucleocytosolic depletion of the energy metabolite acetyl-coenzyme a stimulates autophagy and prolongs lifespan. Cell Metab. 2014; 19: 431–444.

40. Green DR, Galluzzi L, Kroemer G. Metabolic control of cell death. Science. 2014; 345: 1250256.

41. Moussaieff A, Rouleau M, Kitsberg D, Cohen M, Levy G, Barasch D, et al. Glycolysis-mediated changes in acetyl-CoA and histone acetylation control the early differentiation of embryonic stem cells. Cell Metab. 2015; 21: 392–402.

42. Pietrocola F, Galluzzi L, Bravo-San Pedro JM, Madeo F, Kroemer G. Acetyl coenzyme A: a central metabolite and second messenger. Cell Metab. 2015; 2: 805–821.

43. Shi L, Tu BP. Acetyl-CoA induces transcription of the key G1 cyclin *CLN3* to promote entry into the cell division cycle in *Saccharomyces cerevisiae*. Proc Natl Acad Sci U S A. 2013; 110: 7318–7323.

44. Boroughs LK, DeBerardinis RJ. Metabolic pathways promoting cancer cell survival and growth. Nat Cell Biol. 2015; 17: 351–359.

45. Filipp FV, Scott DA, Ronai ZA, Osterman AL, Smith JW. Reverse TCA cycle flux through isocitrate dehydrogenases 1 and 2 is required for lipogenesis in hypoxic melanoma cells. Pigment Cell Melanoma Res. 2012; 25: 375–383.

46. Keller A, Peltzer J, Carpentier G, Horváth I, Oláh J, Duchesnay A, et al. Interactions of enolase isoforms with tubulin and microtubules during myogenesis. Biochim Biophys Acta. 2007; 1770: 919–26.

47. Durrieu C, Bernier-Valentin F, Rousset B. Microtubules bind glyceraldehyde 3-phosphate dehydrogenase and modulate its enzyme activity and quaternary structure. Arch Biochem Biophys. 1987; 252: 32–40.

48. An S, Deng YJ, Tomsho JW, Kyoung M, Benkovic SJ. Microtubule-assisted mechanism for functional metabolic macromolecular complex formation. Proc Natl Acad Sci U S A. 2010; 107: 12872–12876.

49. Noujaim M, Bechstedt S, Wieczorek M, Brouhard GJ. Microtubules accelerate the kinase activity of Aurora-B by a reduction in dimensionality. PLoS One. 2014; 9: e86786.

50. Carbonaro M, O’Brate A, Giannakakou P. Microtubule disruption targets HIF-1alpha mRNA to cytoplasmic P-bodies for translational repression. J Cell Biol. 2011; 192: 83–99.

51. Jaglin XH, Poirier K, Saillour Y, Buhler E, Tian GL, Bahi-Buisson N, et al. Mutations in the β-tubulin gene *TUBB2B*result in asymmetrical polymicrogyria. Nat Genet. 2009; 41: 746–752.

52. Huffaker TC, Thomas JH, Botstein D. Diverse effects of beta-tubulin mutations on microtubule formation and function. J Cell Biol. 1988; 106: 1997–2010.

53. Ti SC, Pamula MC, Howes SC, Duellberg C, Cade NI, Kleiner RE, et al. Mutations in human tubulin proximal to the kinesin-binding site alter dynamic instability at microtubule plus- and minus-ends. Dev Cell. 2016; 37: 72–84.

54. Zhang LG, Li BC, Zhang Y, Jia XJ, Zhou MG. Hexokinase plays a critical role in deoxynivalenol (DON) production and fungal development in *Fusarium graminearum*. Mol Plant Pathol. 2016; 17: 16–28.

55. Bruno KS, Tenjo F, Li, L, Hamer JE, Xu JR. Cellular localization and role of kinase activity of PMK1 in *Magnaporthe grisea*. Eukaryot Cell. 2004; 3: 1525–1532.

56. Zhou YJ, Zhu YY, Li YJ, Duan YB, Zhang RS, Zhou MG. β_1_ tubulin rather than β_2_ tubulin is the preferred binding target for carbendazim in *Fusarium graminearum*. Phytopathology. 2016; 106: 978–85.

57. Yang Q, Yan L, Gu Q, Ma, ZH. The mitogen activated protein kinase BcOs4 is required for vegetative differentiation and pathogenicity in *Botrytis cinerea*. Appl Microbiol Biotechnol. 2012; 96: 481–492.

58. Gu Q, Zhang CQ, Yu FW, Yin YN, Shim WB, Ma ZH. Protein kinase FgSch9 serves as a mediator of the target of rapamycin and high osmolarity glycerol pathways and regulates multiple stress responses and secondary metabolism in *Fusarium graminearum*. Environ Microbiol. 2015; 17: 2661–2676.

59. Duan YB, Xiao XM, Li T, Chen WW, Wang JX, Bart AF, et al. Impact of epoxiconazole on Fusarium head blight control, grain yield and deoxynivalenol accumulation in wheat. Pestic Biochem Physiol. 2018; 152: 138–147.

60. Li J, Duan YB, Bian CH, Pan XY, CJ, Wang JX, et al. Effects of validamycin in controlling Fusarium head blight caused by *Fusarium graminearum*: Inhibition of DON biosynthesis and induction of host resistance. Pestic Biochem Physiol. 2019; 153: 152–160.

61. Livak KJ, Schmittgen TD. Analysis of relative gene expression data using real-time quantitative PCR and the 2(T)(-Delta Delta C) method. Methods. 2001; 25: 402–408.

